# Study on community knowledge and awareness of invasive species

**DOI:** 10.1101/2025.02.18.638815

**Authors:** Leonel Stazione

## Abstract

In a globalized world, invasive species (IS) have significant ecological and socio-economic impacts, emphasizing the need for public awareness and effective management. This work presents a study about environmental perceptions and biological invasion knowledge on both teacher and non-teacher communities, using a semi-structured, online questionnaire. Data on demographic profiles, perceptions of environmental impact and knowledge of biological invasions were compiled. Results revealed that teachers perceived higher environmental and biological invasion impacts than non-teachers. While most respondents acknowledged the high impact of IS, teachers exhibited higher levels of awareness and concern. The unintentional transport of IS was identified as the main threat, and human health concerns were the primary reason for species removal. This study highlights the importance of enhancing environmental education in both formal and non-formal settings to address the impacts of IS. A multidisciplinary approach is recommended to raise public awareness and ensure sustainable management across diverse community sectors.

## Introduction

In this world of constant change and largely connected by globalization the Invasive Species (IS) represent one of the main threat that affect ecosystems and maintenance of biodiversity (Pyšek et al. 2020). The species movement to non-native regions causes profound changes, leading to a process of biotic homogenization between regions (Capinha et al. 2015). When the species are moved far from their native ranges into new regions they can overcome different biogeographical and ecological barriers, with escalating impacts on the environment, economies and social activities (de Melo et al. 2021; Remmele and Lindemann-Matthies 2020). During the last 50 years the transport of IS to non-native regions has intensified and is now considered one of the five main causes of deep global change in nature (IPBES 2023). In addition, the generated impacts for IS problems have important ecological and socioeconomic effects (Bacher et al. 2018), mainly regarding agriculture, forestry and health sectors, generating associated costs for their control and eradication, which directly affect the economies of the countries (Hulme et al. 2024; Latombe et al. 2023).

Considering IS as a serious and growing problem, it is a priority to work on ways to reduce its impacts and avoid new biological invasion processes. In this sense, the public awareness and education are important factors in tackling this issue, since public opinions and attitudes can potentially affect continued introductions and management of IS, it is imperative to understand the public’s level of knowledge and attitudes toward these pests (Kapitza et al. 2019). Previous research on biological invasions has recognized the importance of social perceptions of IS with the majority of studies focusing on the general public (Haley et al. 2023; Lipták et al. 2024; Sosa et al. 2021). However, social perception of IS and attitudes toward their control are controversial, and have conflicts mainly related with biosecurity measures for the multiple users of especially vulnerable ecosystems (Dunn et al. 2018; Shine and Doody 2011; Sutcliffe 2018). Social conflicts seem to be related to the low public knowledge about the correct management of the species and the potential damage it could cause to the ecosystem and to health (Hulbert et al. 2023; Lipták et al. 2024). For that the public education should be considered as an essential part to management of IS.

The education about IS, is especially relevant for the community during their early educational stages - mainly in primary and secondary school-, since children and adolescents often have a poor contact with nature and their attitudes about nature environments are still influenced mainly by Internet content, based on a few iconic and charismatic species, generally exotic. This could interfere with the concept of invasive species and the dangers associated with their establishment (Ballouard 2011; Díez et al. 2018). However, the biological invasions topic is relatively new and it has only recently become relevant to environmental educators (Waliczek et al. 2017). Therefore, knowledge and the optimization of its teaching are constantly changing, which makes the scientific literacy on invasive species among teachers a real determining factor. Teachers are usually seen as mediators between scientists and students. But teaching about biodiversity conservation issues as IS, which involve ecological, economic, and social aspects, presents difficulties for teachers (Borg et al. 2012). Moreover, their personal and professional perception of the environment and IS does have an impact on how teachers approach their tasks as mediators on student learning. The controversial nature of environmental issues and the intrinsically complex and abstract construct of biodiversity might hinder (Büssing et al. 2018).

In order to work on the specific problems of community’s perception of IS, it is essential to determine the knowledge and attitudes of the wider educational and non-educational community about biological invasions, and to identify concepts that can be introduced or reinforced mainly in educational programs.

The general objective of this study is to analyze the perception and knowledge of IS of the educational (teachers) and non-educational (non-teachers) communities of Argentina. In addition, the main objectives are: (1) to explore whether there are differences between the perception of the environment, its biotic components, and global and local threats between the non-educational and the educational communities, as well as within the educational community between teachers of natural sciences (NsT) and teachers of other subjects (non-NsT); and 2) to evaluate whether there are differences on awareness of biological invasions between the non-educational and the educational communities, as well as within the educational community (between NsT and non-NsT).

## Materials and Methods

### Data collection

A survey was designed to query a sample of individuals from this educational (teachers) and non-educational (non-teachers) community regarding their perception of the environment and their understanding of biological invasions. It consisted of an online semi-structured questionnaire of 14 sequenced questions using Google Forms with an introductory paragraph indicating the purpose of the study. The questionnaire covered three main areas: 1-characterisation of the respondents, 2-perception of the environment (biotic components and environmental threats) and 3-awareness of biological invasions (Table 1).

**Table 1.**
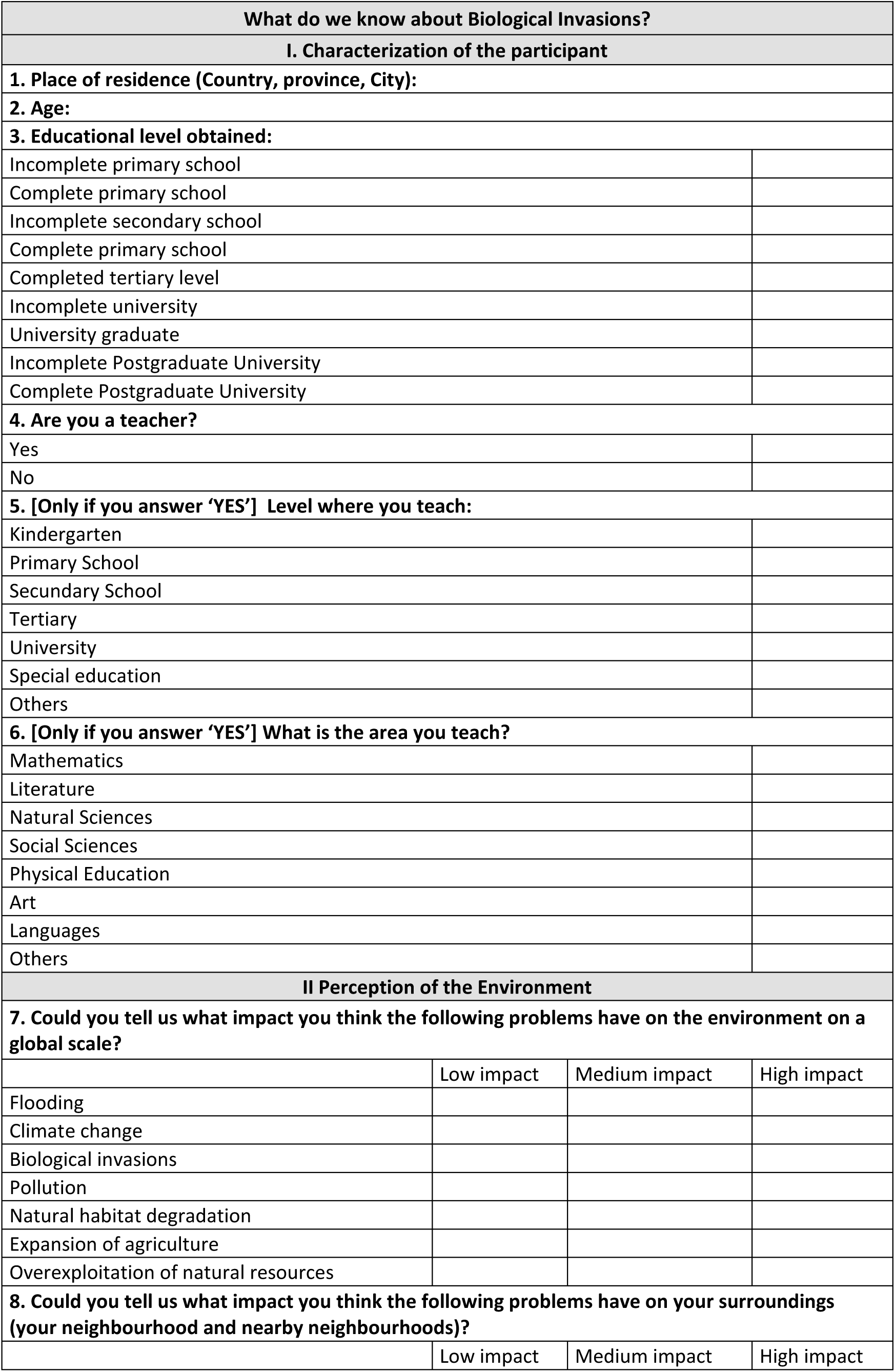
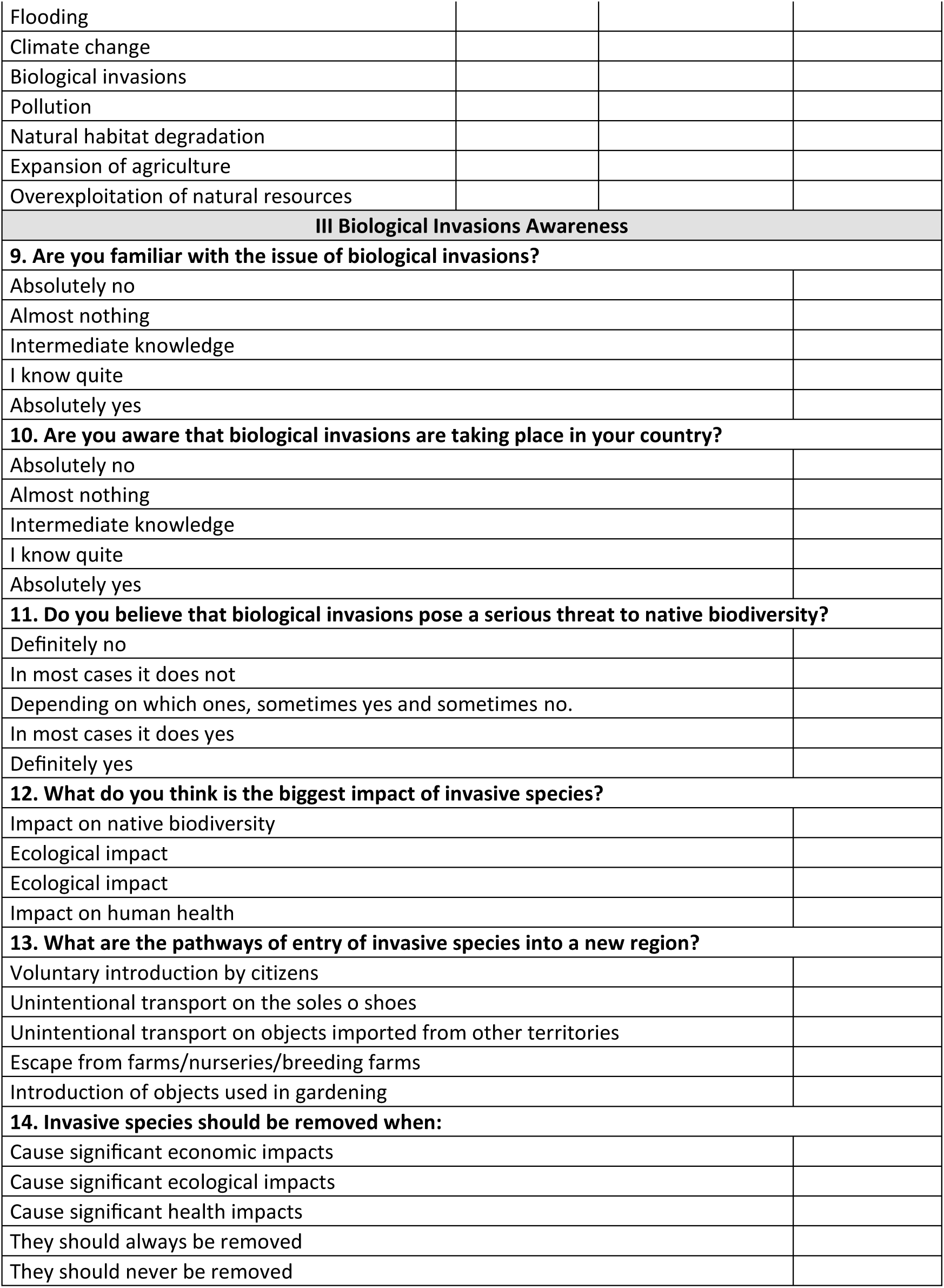
Semi-structured online questionnaire by Google Forms, used to question a sample of individuals from the educational (teachers) and non-educational (non-teachers) community about their perception of the environment and their understanding of biological invasions.

A sampling approach through emails and social media platforms was used. Responses were recorded during 124 days (between June 3 and October 4, 2024). The questionnaire was anonymous in terms of personal information of the participants. The methods were carried out in accordance with the regulations of the Ethics Committee of the Faculty of Exact and Natural Sciences of the University of Buenos Aires, and formal written consent was not required in accordance with the guidelines and regulations, thus confirming the lack non-need for informed consent in the case of an anonymous survey such as the one in this study.

### Characterization of the respondents

To characterize the socio-demographic profile, the respondents were asked to answer: their place of residence, age, education level obtained and if they are related teachers or not. If they were teachers, they were asked: level and subject (Table 1).

### Perception of the environment

In this section, respondents were asked to rate from 1 to 3 (1-Low impact, 2-Medium impact and 3-High impact) the impact of seven threats (flooding, climate change, biological invasion, pollution, natural habitat degradation, expansion of agriculture and overexploitation of natural resources) have on the environment at global and local scale (Q7 and Q8; Table 1). The threats were analyzed by Generalized Linear Model (GLM) analysis using lme4 package in R (version 4.4.1). GLM were performed 1) considered the scale -local and global- as factor, 2) considered Teaching relationship -Teacher or non-Teacher- as factor for each scale (local and global) separately and 3) considered Teaching subjects as factor where two groups were established, one with all teachers teaching natural sciences -NsT- and the other with teachers teaching other subjects -non-NsT-, for each scale (local and global) separately.

### Awareness of biological Invasions

Perception of biological invasions was addressed in six total questions. Respondents were asked whether they were familiar with the issue of biological invasions and if they were aware that biological invasions are taking place in their place (Q9 and Q10). Were also asked about their consideration of invasive species as a serious problem (Q11 and Q12), what would be the main method of entry (Q13) and in which case they consider that IS should be eliminated (Q14; Table 1). To analyze whether there were differences between the different groups of respondents, for each of the six questions separately, the frequency of answers was analyzed and compared using the Chi-square (*X^2^*) test, both between Teaches vs non-Teachers as well as between NsT vs non-NsT.

In addition, to analyze possible correlations between Q9, Q10 and Q11, the questions were ranked from 1 to 5, from “definitely no”=1 to “definitely yes”=5. And Pearson correlation analyses were carried out between Q9, Q10 and Q11 for different participant groups separately: all respondents, Teachers, non-Teachers, NsT, non-NsT. All analyses were performed with R (version 4.4.1).

## Results

A total of 604 responses were spread across the country, with the results covering 22 of the 23 provinces of the country (Figure 1). The participants covered a wide range of ages (mean=39.51 years old, max=73, min=19). According to the level of education attained by the participants, 28% have completed secondary school, 4% have incomplete secondary school, 22% have completed tertiary studies, 22% have a university degree, 16% are currently studying for a university degree, 5% have a completed postgraduate degree and 3% are currently studying for a postgraduate degree.

**Figure 1.**
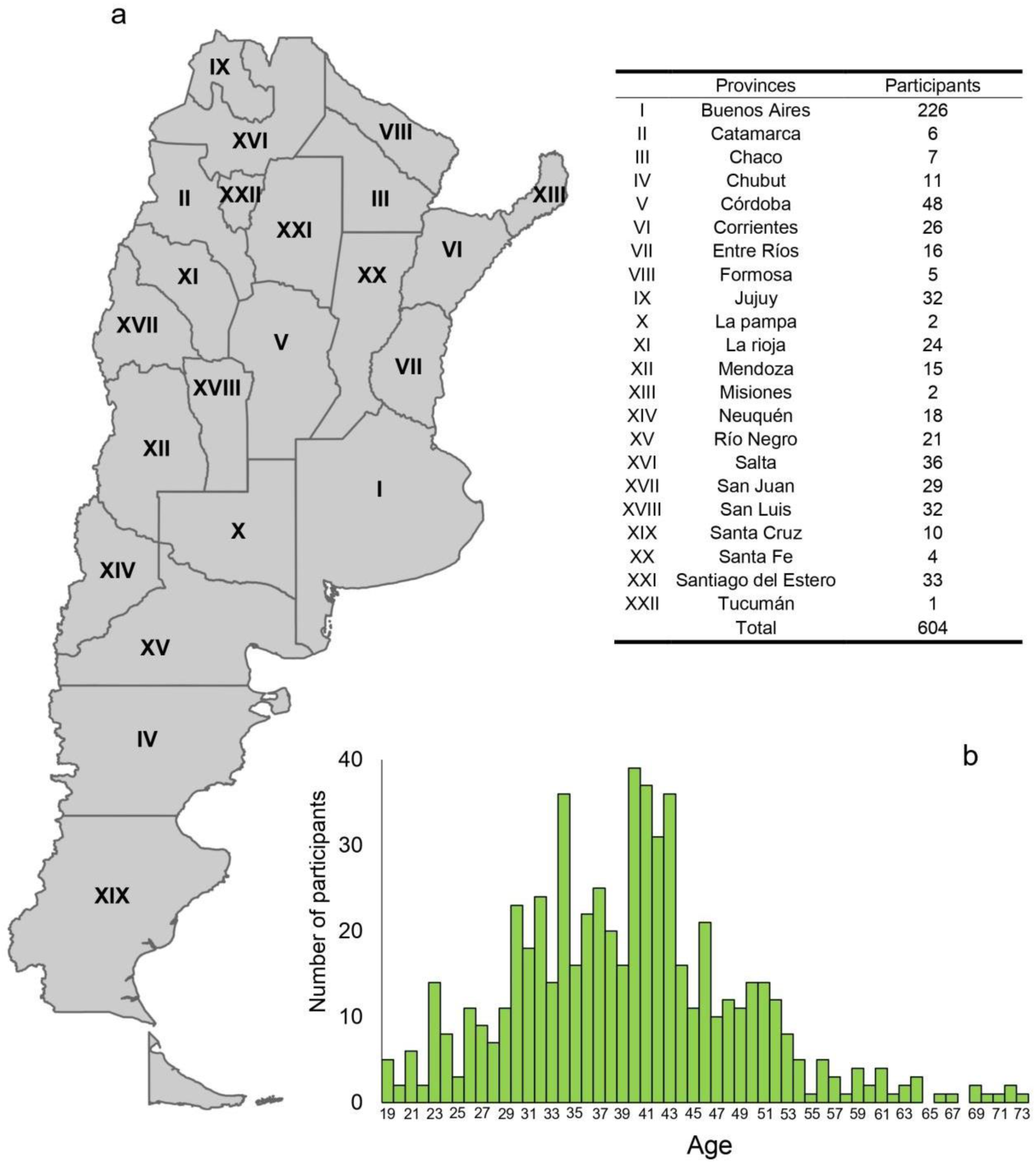
Number of respondents in each Argentine province (a). Histogram of respondents’ ages (b).

Among the 604 participants who filled in the form, 312 were teachers and 292 were no teachers. A total of 27 were kindergarten teachers, 71 primary school teachers, 171 secondary school teachers, 1 tertiary level teachers and 41 university level teachers. Among these, 165 were NsT while 147 were non-NsT.

### Perception of the Environment

The results showed that most respondents considered all threats caused medium to high impact on the environment and that environmental threats -except for flooding-have a significantly higher impact globally than locally (Figure 2a). When analyses were performed for teachers and non-teachers separately, results show that, at both global and local scale, teachers considered that all environmental threats have a significantly higher impact than non-teachers (Figure 2b and 2c). Whereas when the same analyses were conducted separately for NsT and non-NsT, on both global and local scales, NsT considered all environmental threats to have a significantly greater impact than non-NsT (Figure 2d and 2e).

**Figure 2.**
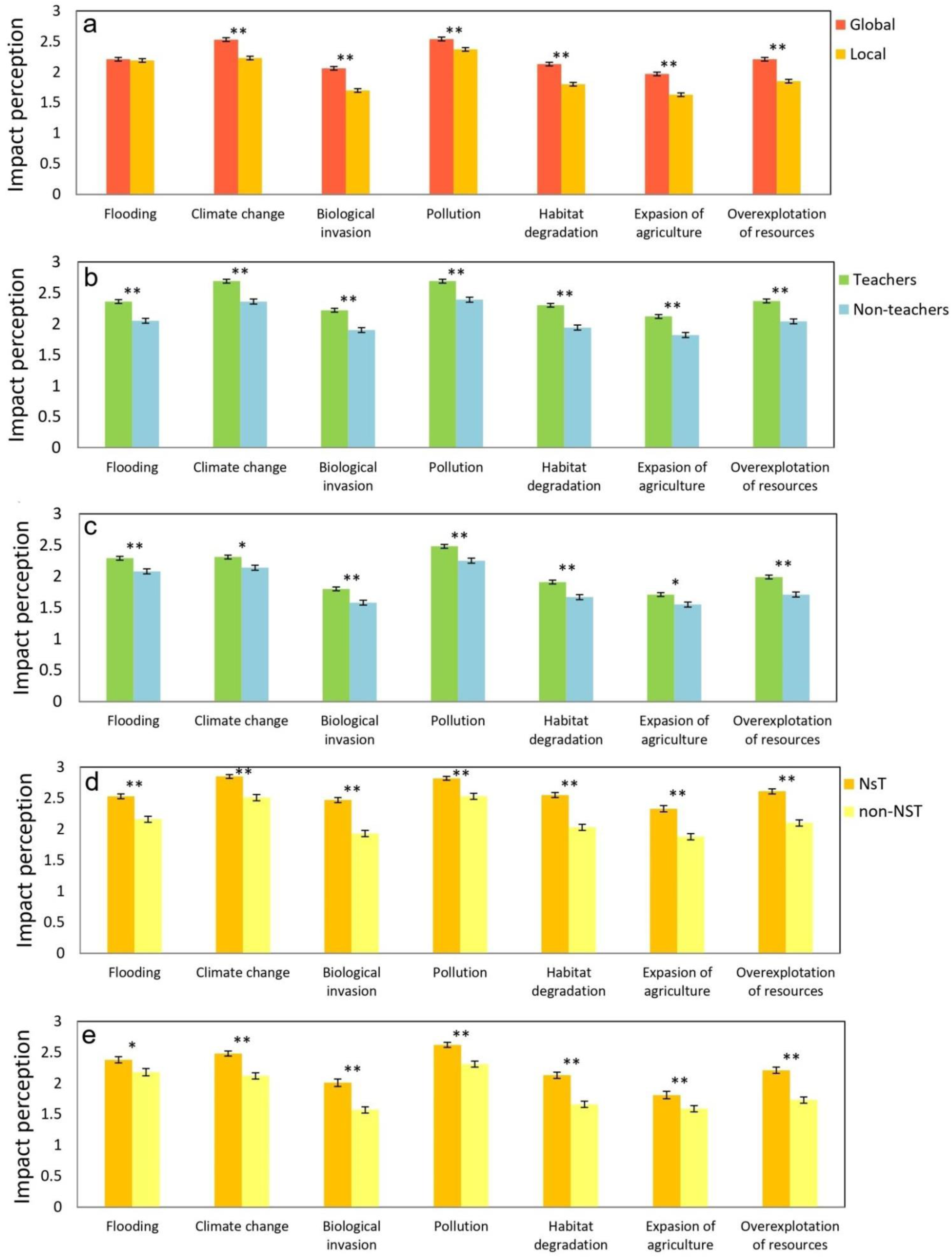
Impact perception of different threats to the environment: at global and local scale (a); for Teachers and Non-teachers at global (b) and local (c) scale; and for NsT and non-NsT at global (d) and local (e) scale. Asterisks indicate significant differences (*p < 0.01; **p < 0.001).

### Awareness of biological Invasions

The results show that 43% of respondents overall reported knowing almost nothing about biological invasions (Q9), while that 33% reported intermediate knowledge. But when teachers and non-teachers are analyzed separately, there are significant differences in the answers given between the two groups (*X^2^_df=4_*=94.5, p<0.001). Most teachers responded with intermediate knowledge (51%) while most non-teachers responded that they know almost nothing (66%, Figure 3a). Among teachers, the NsT showed higher knowledge about biological invasions than non-NsT (*X^2^_df=4_*=, p<0.001, Figure 3a). Regarding to Q10 the most reported low knowledge about current biological invasions. But when teachers and non-teachers are separated, the results show significant differences where teachers have more knowledge about the problem than non-teachers (*X^2^_df=4_*=90.02, p<0.001, Figure 3b). Also differences where founded between NsT and non-NsT (*X^2^_df=4_*=94.3, p<0.001), the most of NsT declared an intermediate knowledge (50%) while that the most of non-NsT responded that they know almost nothing (52%, Figure 3b).

**Figure 3.**
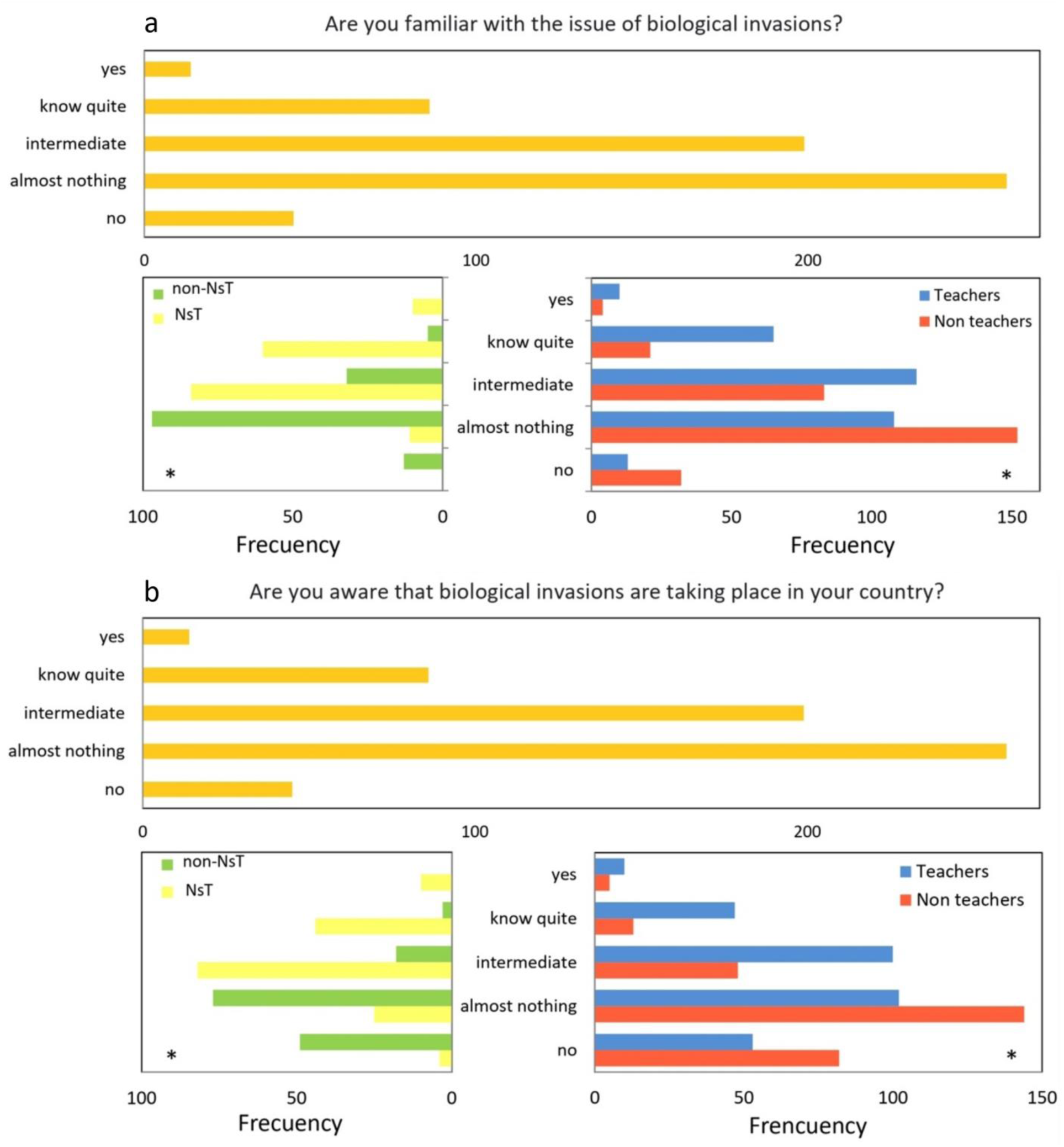
Knowledge of biological invasions -Q9- (a) and Awareness of biological invasions -Q10- (b) from: all respondents and also partitioned between Teachers and Non-teachers; and Nst and non-NsT. Asterisks indicate significant differences.

For the Q11 about whether biological invasions pose a serious risk to native biodiversity, the most of the answers (42%) were “sometimes yes/sometimes no”. Differences were found in the answers between teachers and non-teachers (*X^2^_df=4_*=93.4, p<0.001) as well as between NsT and non-NsT (*X^2^_df=4_*=57,5, p<0.001, Figure 4a). Regarding the impact of biological invasions (Q12), the most frequent responses were impact on native biodiversity (35%) and on human health (29%). Results showed significant differences between teachers and non-teacher responses (*X^2^_df=4_*=66.7, p<0.001) where the most frequent responses from the teacher was native biodiversity (49%) while from non-teachers was human health (38%, Figure 4b). Also, analyses showed significant differences on responses between NsT and non-NsT (*X^2^_df=4_*=72.4, p<0.001) where the most frequent response from NsT was native biodiversity (73%) and from non-NsT was human health (39%, Figure 4b).

**Figure 4.**
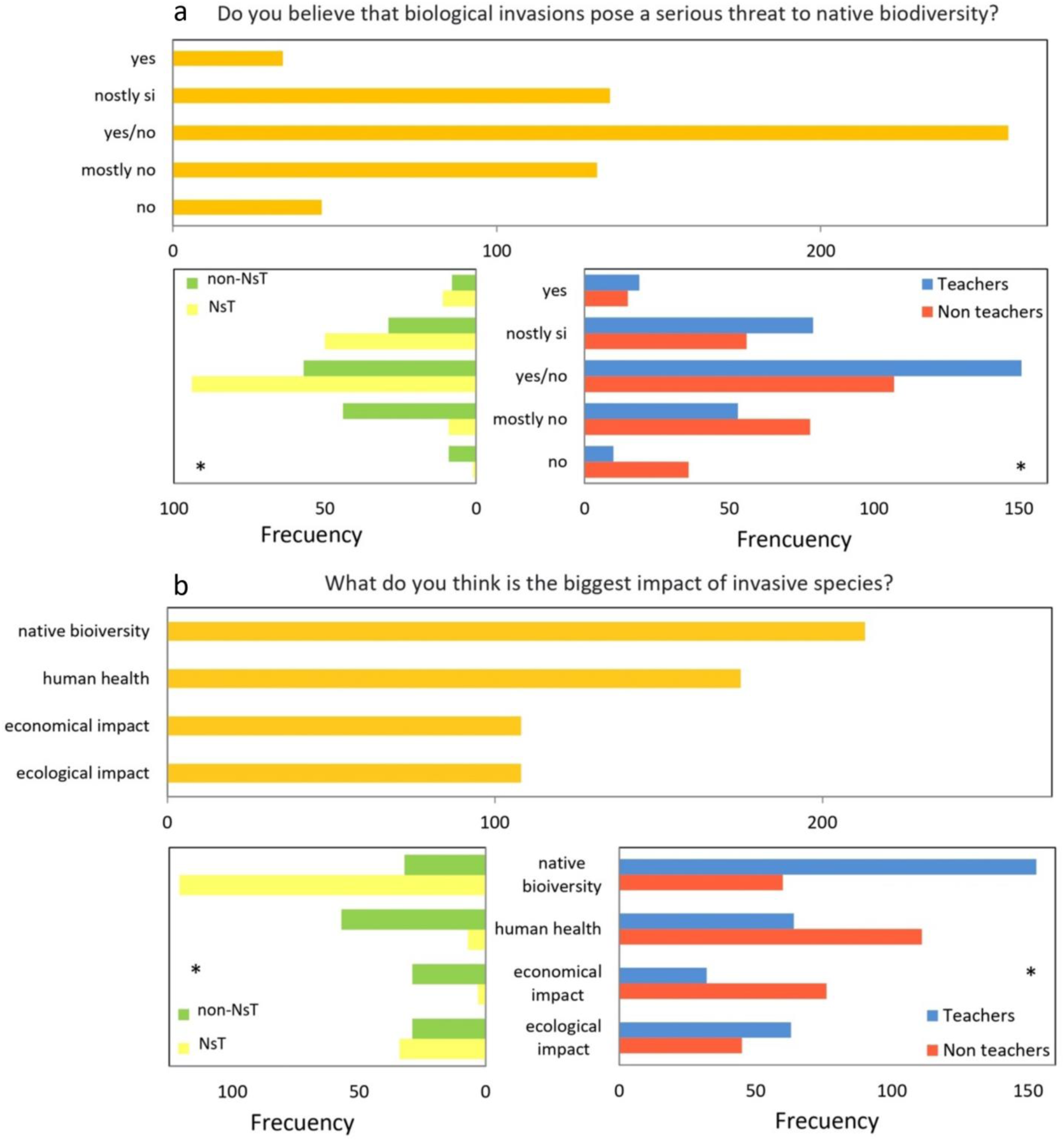
Perception of the level of impact -Q11- (a) and perception of type of impact of biological invasions -Q12- (b) from: all respondents and also partitioned between Teachers and Non-teachers; and Nst and non-NsT. Asterisks indicate significant differences.

For the Q13 about entry pathways of invasive species, the results show that the majority of respondents answered that the most important route is unintentional transport on imported objects (51%, Figure 5a). Differences were found in the answers between teachers and non-teachers (*X^2^_df=4_*=53.4, p<0.001) as well as between NsT and non-NsT (*X^2^_df=4_*=48.5, p<0.001). However, in all cases the response patterns are similar, with “unintentional transport on imported objects” being the most frequent response (Figure 5a). In Q14 on when invasive species should be removed, the most frequent answer was when it affected human health (48.7%). Analysis showed differences between teachers and non-teachers (*X^2^* =79.3, p<0.001), where the most frequent answer from both groups was human health impacts. Also differences were found between NsT and non-NsT (*X^2^* =89.4, p<0.001), where the most frequent response from NsT was ecological impacts (45%) and from non-NsT was human health impacts (64%, Figure 5b).

**Figure 5.**
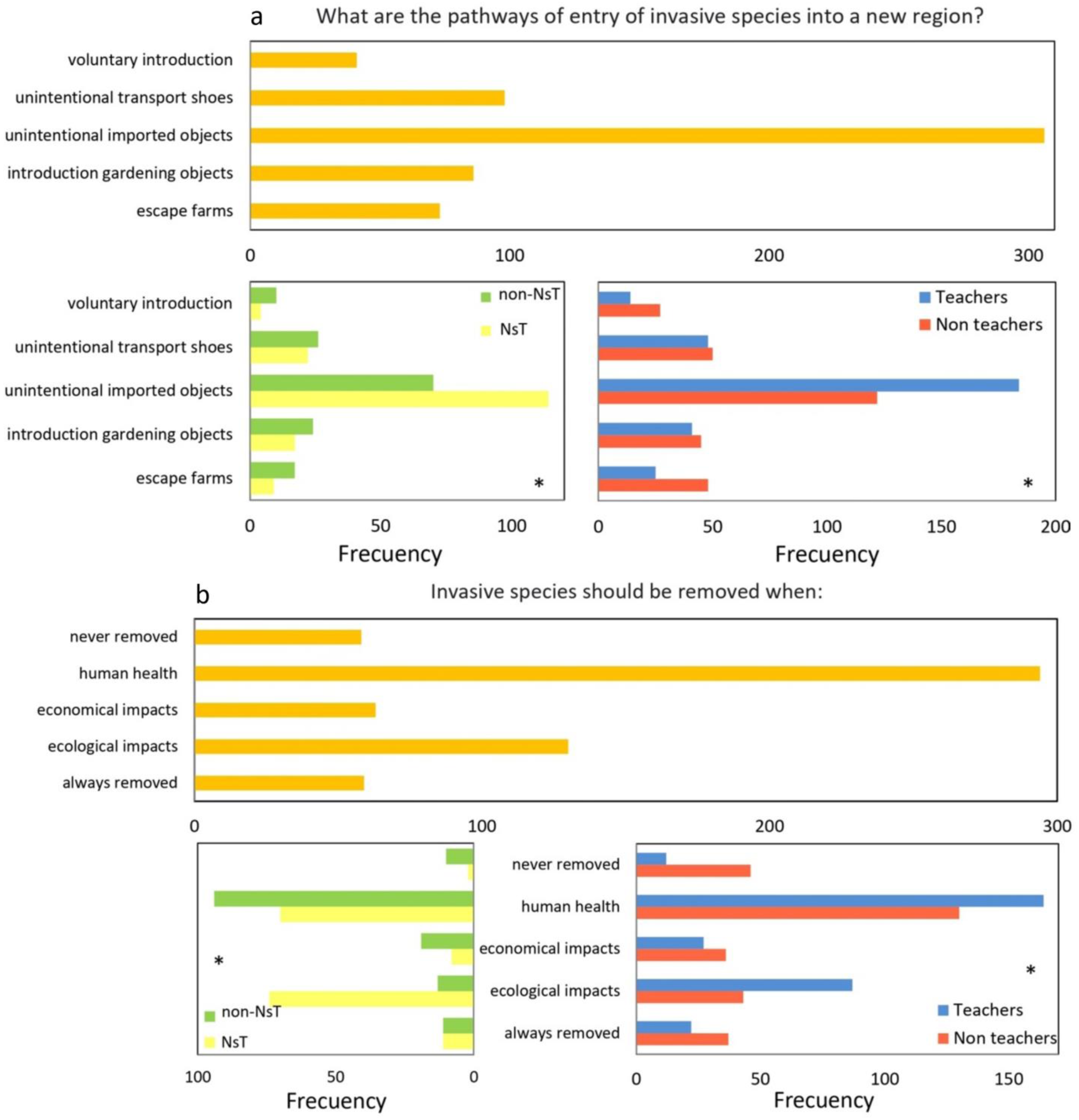
Perception of the main pathways of entry of IS -Q13- (a) and responses about when IS should be removed -Q14- (b) from: all respondents and also partitioned between Teachers and Non-teachers; and Nst and non-NsT. Asterisks indicate significant differences. Asterisks indicate significant differences.

The Pearson correlations between Q9 and Q10 were significant in all cases. While both between Q9 and Q11 and between Q10 and Q11 were significant for all cases except non-teachers (Table 2).

**Table 2.** Pearson correlation coefficients between Q9, Q10 and Q11. For all respondents together, as well as for the groups of teachers, non-teachers, NsT and non-NsT, separately. Asterisks indicate statistically significant correlations (**p < 0.01; ***p < 0.001).

| Variables |  | Participant groups |  |  |  |  |
| --- | --- | --- | --- | --- | --- | --- |
| *V1* | *V2* | All | Teachers | non-Teachers | NsT | non-NsT |
| *Q9* | *Q10* | 0.77*** | 0.83*** | 0.67*** | 0.08*** | 0.56*** |
| *Q9* | *Q11* | 0.28*** | 0.42*** | 0.08 | 0.22** | 0.43*** |
| *Q10* | *Q11* | 0.26*** | 0.40*** | 0.06 | 0.26*** | 0.35*** |

## Discussion

With the increasing environmental degradation of ecosystems, more sustainable environmental behavior and management is necessary for the sustainable maintenance of the environment (Olsen et al. 2020). To this end, it is important to ensure that the community -mainly through education-is provided with adequate knowledge to enable the acquisition of optimal skills and attitudes (Parra et al. 2020). In addition to greater knowledge of ecological concepts and processes that provide the basis for understanding the human impact on the functioning of ecosystems and ecosystem services, such as IS, which represent one of the main threats to biodiversity worldwide (Gallardo et al. 2019).

According to the data obtained, the study shows that the community in general perceives that the impact of the different threats to the environment to be greater at the global level than at the local level, in most of the factors involved in the survey. These factors were also considered the most damaging in other studies related to the educational environment (Sosa et al. 2021; Vilches et al. 2015), alluding responsibility to demographic growth of the human population. Also, the results indicate that the educational community shows a greater impact perception and concern for environmental drivers than the non-educational community. Despite the fact that biological invasions represent one of the main drivers of biodiversity loss, this work showed that both educational and non-educational communities considered it less important both globally and locally than other drivers. Similar results have also been reported in previous studies (Bermudez et al. 2020; Sosa et al. 2021; Vilches et al. 2015), which could indicate the absence of biological invasion issues in educational curricula that representatively reflect the threat that IS pose to the environment and the ecological effects they can have (Remmele and Lindemann-Matthies 2020; Waliczek et al. 2017).

When investigating the knowledge of biological invasions and the impact they have on native biodiversity the results showed that teachers were more aware of biological invasions -especially NsT-than non-teachers. In addition, a positive correlation was observed between the level of knowledge about IS and the effect they have on native biodiversity in all groups of teachers, while this correlation was not found in the non-teaching community. Considering that the biological invasions will continue to increase in the next decades, the impacting on biodiversity, ecology, and socio-economic demanding urgent management policies (Busso et al. 2013; Seebens et al. 2021); and the optimal reception and acceptance of IS management -involving all citizens-could be related to having a background of environmental knowledge (Ekanayake et al. 2020; Waliczek et al. 2017). This study shows differences in the current background of the educational community compared to the general community (non-teachers) which could help to obtain greater support and compromise in the inclusion of most topics related to biological invasion in different educational levels, involving to educators and non-educators through the inclusion of transversal themes in all educational curricula (Gayford 2014; Verbrugge et al. 2021).

Regarding the question about the biggest impacts caused by IS, differences were found between the groups, where NsT identified the native biodiversity impact as the highest problems while the non-teachers and non-NsT identified human health impacts as the greatest problems. These differences suggest that perception and awareness of environmental effects may depend on the environmental and ecological knowledge of the community (Kapitza et al. 2019). However, both native biodiversity and human health impacts highlighted over other options, pointing to the socio-ecological problem of IS (Kannan et al. 2014; Kelsch et al. 2020). On other hands, participants -both teachers and non-teachers-have widely recognized the “unintentional imported objects” as the main pathway to the IS. However, they had difficulties recognizing different IS pathways to entry, especially the transport on the soles of shoes and other personal objects, or the escape from farms. As for the lack of awareness of unintentional transport of personal objects, this could indicate that respondents are not aware of their own role as potential vectors of IS. This unawareness is especially relevant in protected areas or national parks where visitors can act as vectors for the transport of spacers (Ford-Thompson et al. 2015). These results, together with previous evidence, highlight the relevance of an appropriate education, either formal or informal, about IS (Ladrera et al. 2020; Remmele and Lindemann-Matthies 2020). This education should lead the educational and non-educational community to understand IS potential threat to biodiversity and the negative impact on the appropriate functioning of the whole socio-ecological system, including economic and also human health threats.

The majority of respondents considered that invasive species should be removed especially when they cause human health problems. In the case of NsT, secondly they considered the need to remove invasive species when they cause ecological impacts. But in all cases, a very small proportion of respondents pointed to the need to always remove IS from invaded territories. These results indicate that the respondents are not aware that all IS, by definition, cause impacts of a different nature and, therefore, their elimination implies benefits of different types. Therefore, these results highlight the need for the community in general to understand the reasons for carrying out certain IS management strategies. Ladrera et al (2020) showed that the human consideration of IS as potential danger factors depend on the species since human beings tend to empathize more with species similar to us, fundamentally mammals, for which greater care and conservation efforts are exercised. Whereas there is a greater social rejection of other types of organisms, mainly invertebrates, which could favor greater support for the sacrifice of invasive taxa of this type (Leandro et al. 2019). The social rejection of certain IS control measures has been extensively studied and several works have pointed to the importance of robust knowledge about biological invasions and environmental issues to favor the development of positive attitudes toward their control (Green et al. 2016; Waliczek et al. 2017). In this sense, more training in the area both within formal and non-formal education -e.g. activities of outreach, awareness rising-could lead to higher knowledge and a better understanding of the problem. Likewise, It can translate into attitudes more inclined toward IS control.

It is urgent to increase the background knowledge of the community on this subject. For that, this work suggests that the knowledge about IS and their potential threats should be focused on educational actors -both teachers and students-as well as citizens outside formal education. In the context of formal education, curricula should promptly acquire changes at all educational levels focusing on a cross-curricular approach to all subjects (Ladrera et al. 2020). Simultaneously, in the non-formal education context the outreach activities in strategic establishments such as museums, nature parks, ecological reserves, gardens and eco-parks could represent fundamental tools to increase and disseminate environmental knowledge and awareness to the general community (Reed et al. 2010). On other hands, IS knowledge needs to be addressed in a multidisciplinary approach and in a variety of ways, as environmental problems involve and are related to many social problems such as, health, ecology, economy, ancient culture, religions, etc (Estévez et al. 2015).

In the face of the world’s current problems, the ultimate goal is to reinforce public environmental awareness of the role of humans as potential vectors of IS and the benefits associated with the development of control management. This requires the equal inclusion of different sectors of the community through an articulation between politicians, scientists, teachers, students and the wider community.

## Acknowledgements

The author would like to thank all the teachers and non-teachers who accepted the collaboration and completed the form. The author also thanks the editor and the reviewers for their help to improve this article.

## Disclosure statement

No potential conflict of interest was reported by the author(s).

## Notes

### Competing Interest Statement

The authors have declared no competing interest.

